# Do AI Models for Protein Structure Prediction Get Electrostatics Right?

**DOI:** 10.64898/2026.03.11.711144

**Authors:** George I. Makhatadze

## Abstract

A variant of the U1A protein containing four substitutions to ionizable residues was generated serendipitously due to a miscommunication. Biophysical measurements show that this variant has at least twice as much helical structure as the wild-type U1A and is trimeric in solution, in contrast to the monomeric wild type. In sharp contrast, structures predicted by deep-learning AI tools (AlphaFold2 and RoseTTAFold2) and transformer-based tools (OmegaFold and ESMFold) are all highly similar to the wild-type U1A (backbone RMSD < 1 Å). Even more surprising, two of the substituted ionizable residues are predicted to be fully buried in the non-polar core of the protein, an outcome that contradicts well-established physico-chemical principles, as ionizable residues are normally located on the protein surface. To explore this effect further, we generated sequences containing up to all twelve residues that make up the non-polar core of U1A. Across thousands of sequences, and depending on the AI model used, the majority of predicted structures contained fully buried ionizable residues while still maintaining the overall U1A fold. We then examined two additional proteins of comparable size, acylphosphatase and the de novo-designed TOP7 fold, and observed the same phenomenon: AI models frequently predicted structures with buried ionizable residues that nevertheless retained the parent fold. When these AI-predicted structures were subjected to short (50 ns) molecular dynamics simulations using physics-based force fields such as CHARMM or AMBER, the structures rapidly relaxed into ensembles that exposed ionizable residues. We conclude that while AI-based structure prediction tools perform extremely well on naturally occurring sequences, they do not reliably encode the physico-chemical principles governing the placement of ionizable residues. A straightforward remedy is to include a brief molecular dynamics simulation as a final validation step for AI-generated structures.

## Introduction

Deep learning-based protein structure prediction has transformed structural biology over the past several years, reshaping how researchers approach everything from basic mechanistic questions to drug discovery and protein design. The most prominent example, AlphaFold2 (1), demonstrated that many single-chain protein structures can be predicted with near-experimental accuracy directly from sequence, a milestone widely described as transformative for the field and recognized by the 2024 Nobel Prize in Chemistry (2). This breakthrough has been followed by a rapidly growing family of AI-based structure prediction methods (e.g., RoseTTAFold (3), ESMFold (4), OmegaFold (5), and others (6, 7)) and large public structure resources such as the AlphaFold Protein Structure Database (AFDB), which now contains predicted structures for hundreds of millions of proteins (8). At the same time, a growing body of work has begun to systematically probe where these models fail, highlighting important limitations in the ability to predict conformational heterogeneity and functional dynamics (9–14), show overreliance on “memorization” (15–17), and are frequently unable to capture correct biomolecular energetics (18–20). Instances in which predictions produced incorrect and/or incomplete structures were also documented (21–24). Here we provide a compelling example in which AI-based models predict structures that violate known principles of protein structure formation, namely the overwhelming tendency that non-membrane proteins tend to keep the ionizable residues on the protein surface due to highly unfavorable energetics of burying polar or charged groups in the non-polar protein interior (25–32).

The paper is arranged as follows. We begin by presenting experimental structural data, obtained serendipitously, on the effects of four amino acid substitutions on the structure of a small globular protein. We then show that current AI-based protein structure prediction fails to capture large conformational changes arising from these few amino acid substitutions. Moreover, the predicted structures violate basic thermodynamic principles of protein structural organization by burying ionizable residues within the protein’s nonpolar core. We then assess the generality of these effects by analyzing thousands of AI-generated structural predictions for sequences in which we introduced ionizable residues at multiple positions within the hydrophobic cores of three different proteins. This analysis shows that AI-based structure predictions violate the common physico-chemical organizing principles of protein structure and often produce structures with multiple buried ionizable residues. Finally, we show that physics-based force fields (CHARMM or AMBER) can detect these deficiencies in structural predictions within a few picoseconds in all-atom explicit-solvent molecular dynamics simulations.

## Results and Discussion

*“The most exciting phrase to hear in science, the one that heralds new discoveries, is not’Eureka!’ but’That’s **funny**…’”* <u>Isaac Asimov</u>

### Serendipitous Experimental Results on Consequences of Burying Ionizable Residues in U1A protein

A year before the publication of a paper describing the work on increasing protein stability by optimizing interactions between ionizable groups on protein surfaces (33), a curious result was obtained in the lab. A variant of U1A was made in which, due to the mix-up in the residue numbering by 1 between the PDB structure and the expression construct, four ionizable residues were substituted into positions (I14E, G38E, T66E, I84K) preceding the intended ones (N14E, Q39E, N67E, Q85K). Needless to say, the intended substitutions led to the desired stabilization, as was reported in (33). The unintended variant was expressed and purified without major difficulty but exhibited properties dramatically different from those of the wild-type U1A. First, the circular dichroism spectrum of this U1A variant showed a marked difference (**Figure 1A**). The far-UV spectrum of the variant showed a dramatic increase in helical content relative to the wild-type protein: the ellipticity at 222 nm, which is usually directly related to the helical content, is doubled. The addition of 6 M GdnCl, as expected, completely abolishes the CD signal at 222 nm and produces overlapping spectra. The near-UV CD spectra, which reflect the environment of aromatic residues (there are 4 Tyr and 7 Phe residues in the U1A sequence), are rather unremarkable, with slightly higher amplitude for the U1A variant (**Figure 1B**). Again, the addition of 6 M GdnCl leads to a significant loss of signal, producing overlapping spectra for the variant and wild-type U1A. Importantly, equilibrium analytical ultracentrifugation (**Figure 1C**) shows that while the wild-type U1A is monomeric in solution (obtained molecular mass M_m_=13,500±500 Da; expected 12,578 Da), the U1A variant forms a trimer (obtained M_m_=30,600±500 Da; expected 12,709 Da). Taken together with NMR data (see supplementary **Figure S1**), this led to the designation of this U1A variant as “funny-U1A”.

**Figure 1.**
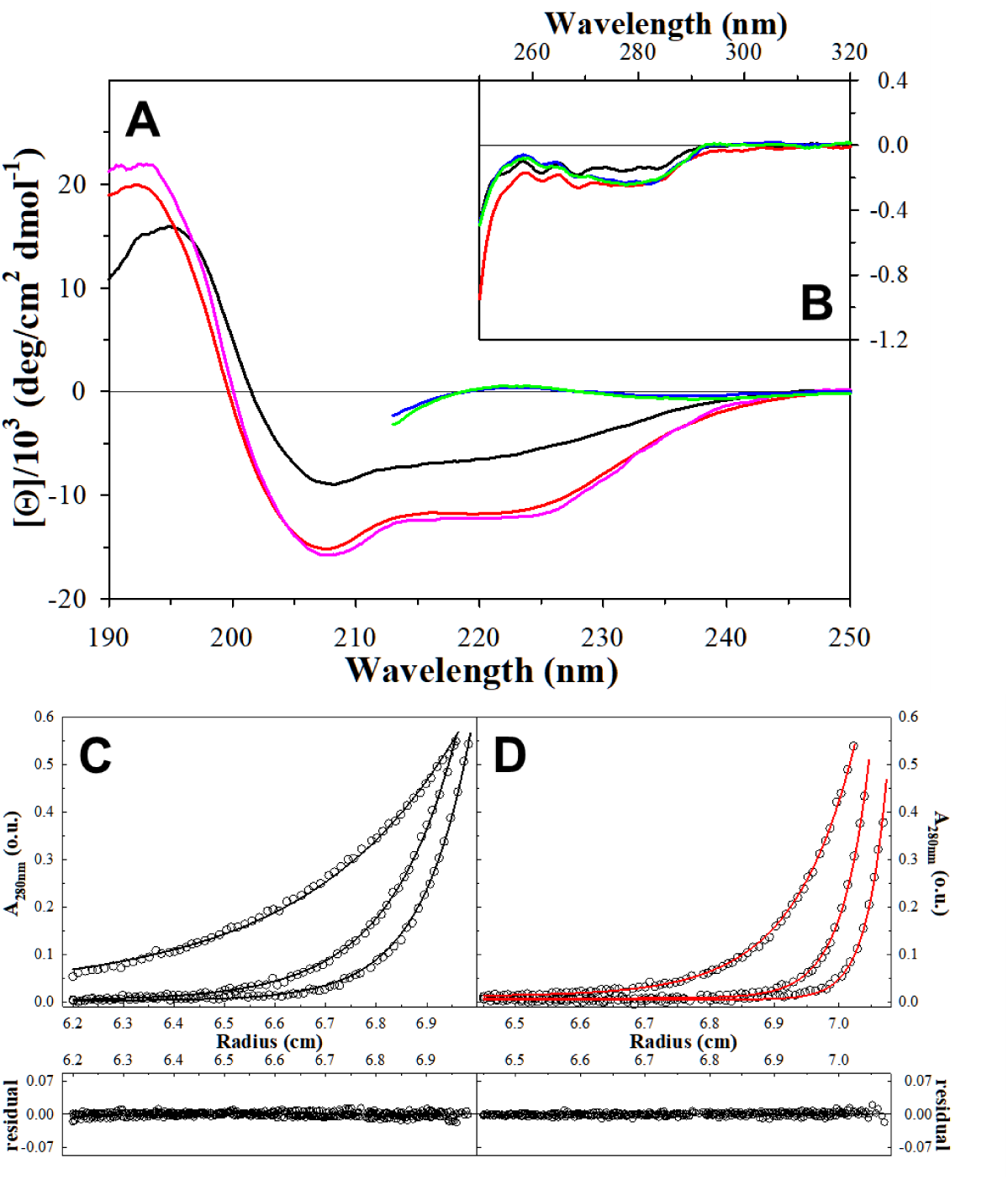
**Experimental Data for U1A proteins. *Panels A-B.*** Comparison of the circular dichroism spectra for the U1A protein variants in the Far-UV (A) and near-UV (B). Black (wt-U1A) and red or pink (funny-U1A) in aqueous buffer; blue wt-U1A and green (funny-U1A) in the presence of 6 M GdnCl. ***Panels C-D.*** Analytical centrifugation profiles of wt-U1AWT (Panel C) and funny-U1A (Panel D). Symbols show the experimental data points (every fifth point is shown for clarity) at three different speeds: 20,000, 30,000, and 35,000 rpm. The solid lines are the fits to equation 1 (Panel C: M_m_=13,500±500 Da; expected 12,578 Da, Panel D: M_m_=30,600±500 Da; expected 12,709 Da). The plots of the residual distributions of the fits are also shown.

Placement of four ionizable residues at the unintended positions in U1A can have several possible consequences. Gly38 is located in the Ccap position of the first α-helix. The interactions at the C-terminus of this helix represent a classical capping box (34) and include a hydrophobic contact between Phe34 and Ile40 (35). Substitutions at Gly at the Ccap position usually decrease protein stability (36–38). Thr66 is located in the middle of the second α-helix (residues 62-72). According to helix-forming propensities, largely defined by the side chain entropy, substitution with Glu in the middle of a helix should increase stability because gamma-branched amino acids have a higher helix-forming propensity than the beta-branched Thr (39). Furthermore, the *i* to *i+4* salt bridge between Glu66 and Arg70 might somewhat increase the stability of this helical segment (40). Two positions, I14 and I84, are located at the adjacent positions across antiparallel β1 and β4 strands. The non-polar side chains of these two residues are fully buried in the core of the U1A protein. Substitutions to ionizable residues at these positions are expected to dramatically decrease stability and should lead to an unfolded protein. And yet, the funny-U1A is folded to an alternate structure. Unfortunately, attempts to crystallize funny-U1A failed, and thus, without a 3D model, the results were not very “publishable” at the time.

### AI-based Prediction of the Effects of Substitutions on the Structure of U1A Variant

Recent breakthroughs in AI-based structural prediction methods have revolutionized biology. The two main flavors of the prediction software differ significantly in their underlying AI type, core architecture, input focus, and training strategy. AlphaFold and RoseTTAfold use deep learning (DL) with attention-based neural networks that combine sequence, structure, and multiple sequence alignment (MSA) data and are trained on evolutionary data and structural databases. In contrast, ESMFold and OmegaFold are transformer-based protein language models (TR) trained on massive protein sequence datasets and use single protein sequences (i.e., they do not require MSA). We applied these four AI-based models to predict the structure of the funny-U1A variant.

The structure of the funny-U1A variant predicted with AlphaFold2 is compared with the PDB structure of the wild-type in **Figure 2**. Contrary to expectations, AlphaFold predicted a structure that was identical to that of the wild-type protein (1.03 Å backbone RMSD from the x-ray structure and 0.47 Å RMSD from the AlphaFold predicted wild-type structure). This contradicts the results of the experimental studies, suggesting large conformational changes in funny-U1A. Running the sequence with the multimer option that forces formation of a trimer produces a structure with an individual fold for each monomer, similar to that of the wild-type protein. What is even more alarming is that the side chains of ionizable residues that are substituted for Ile at positions 14 and 84 remain buried in the nonpolar core of the protein. Moreover, analysis of the multiple sequence alignment (MSA) for U1A indicates that all four substituted positions are highly conserved. Moreover, never or very rarely are these positions occupied by ionizable residues (see **Table 1**). Thus, there is no evidence that AlphaFold predictions of buried ionizable residues at positions 14 and 84 in funny-U1A are guided by the signal in MSA. Importantly, similar results are observed for the structure of U1A variant predicted by other AI-based algorithms such as RoseTTAfold, OmegaFold, and ESMFold (data not shown), again suggesting a common issue for all, DL- or TR-based, AI structure prediction algorithms.

**Figure 2.**
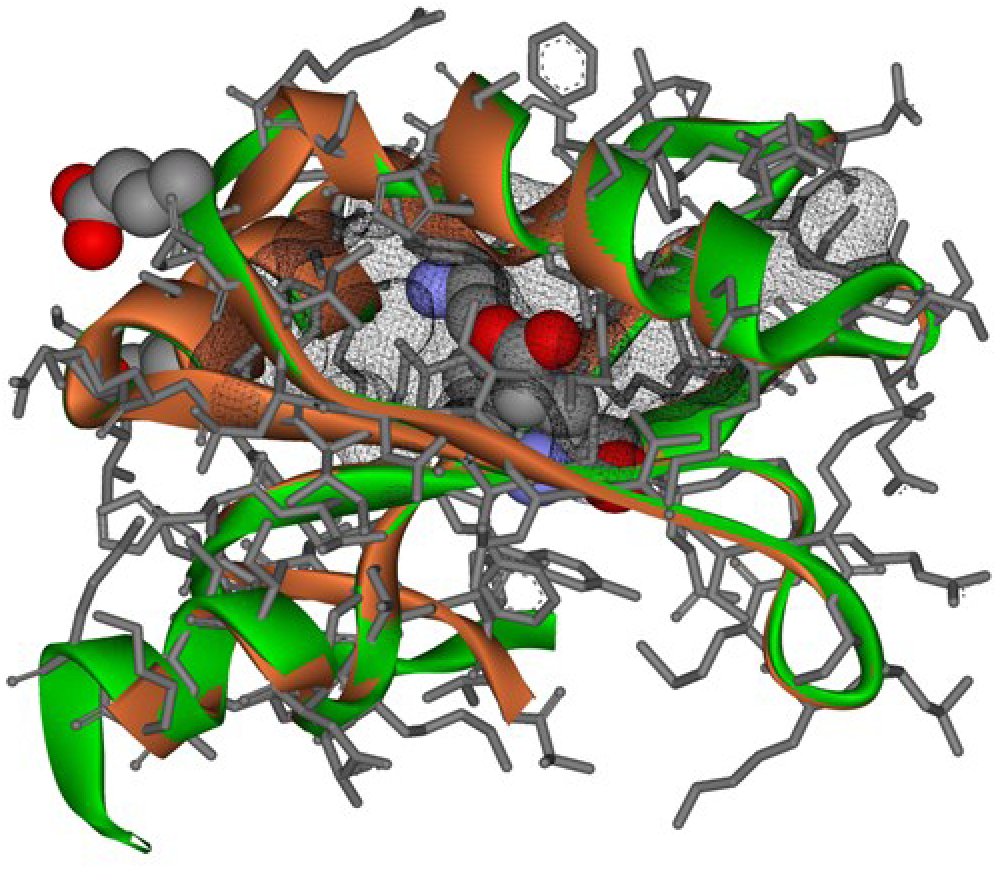
A cartoon of an overlay of the experimental structure of U1A (PDB:1URN, orange cartoon, gray sticks for bonds) with the structure of funny-U1A predicted by AlphaFold2 (green). The non-polar core is shown as gray mesh, and the sites with the substitutions in funny-U1A are shown in CPK representation.

**Table 1.**
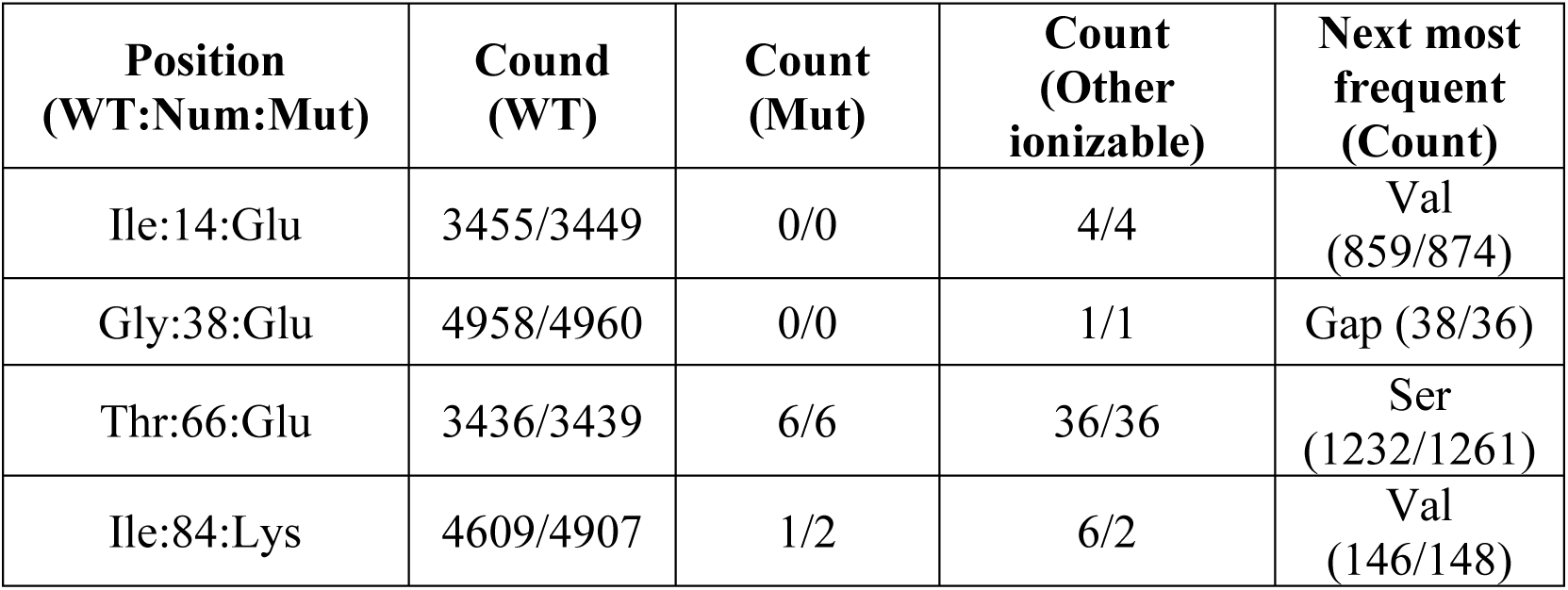
Statistical analysis of frequencies of different residues in the 5001 homologous sequences at the positions corresponding to Ile14, Gly38, Thr66, and Ile84 of human U1A.

Previous statistical analyses of experimentally determined protein structures from the PDB suggest that buried ionizable residues in proteins are either hydrogen-bonded or form salt bridges (27). There is no evidence that buried ionizable residues at positions 14 and 84 of funny-U1A form hydrogen bonds or salt-bridge interactions in any of the AI-predicted structures. It is important to note that studies of the effects of substitutions of non-polar residues with polar residues in the hydrophobic core of the protein show significant destabilization on the order of several kcal/mol (41, 42). The only known systematic studies in which an actually buried ionizable residue was only possible to observe in the context of the native protein were in the case of a hyper-stabilized variant of SNase ΔPHS ΔG=12-14 kcal/mol (43–46).

### How Many Ionizable Residues Can AI-algorithms Tolerate in the Core of a Protein

The predicted structure of the funny-U1A variant with buried ionizable residues might be caused by the fact that having so few substitutions does not yield a significant signal to warrant exploring alternative structures. The rarity of ionizable residues in MSA at substitution positions further exaggerates this (see **Table 1**). To probe the threshold of how many substitutions with ionizable residues in the hydrophobic core of U1A are needed for AI-tools to start exploring alternate structures, we replaced them with different sets of ionizable residues at twelve positions in the non-polar core of U1A. These twelve positions (see **Table S1**) show that according to the analysis of MSA, they are largely (>93%) occupied by hydrophobic residues (defined as Ala, Val, Leu, Ile, Phe, Trp, Tyr) and the fraction of ionizable residues at these positions is less than 0.14%. We used sequence substitutions with Asp, Glu, Lys, and Arg residues across multiple sets of varying numbers of substitutions (1 to 12) at randomly selected sites among 12 core positions, to generate expected structures using AlphaFold2, OmegaFold, and ESMFold. In addition, RoseTTAfold2 was used to generate the expected structure for sequences in which all 12 sites were replaced by one of the four ionizable residues. The resulting structures were analyzed to extract four structural descriptors (see Materials and Methods section for details).

The first descriptor is the average exposed fraction of the surface-accessible area of the side chains of ionizable residues, calculated per residue type (Asp, Glu, Lys, or Arg). This descriptor indicates whether the ionizable residues in the structures are buried within the protein core or exposed to the solvent. The second descriptor is the root-mean-square deviation of the polypeptide backbone from the experimentally determined X-ray structure of the wild-type protein, RMSD (Å). This parameter provides a detailed measure of the similarity between the predicted structures and the PDB structure. The third descriptor is the radius of gyration, Rg (Å), which characterizes the overall compactness of the predicted structure and can be compared with the experimentally determined X-ray structure of the wild-type protein. The fourth descriptor represents the average structure prediction accuracy, as reported by the pLDDT score.

Figure 3 shows the dependencies of these four descriptors on the number of substitutions with the ionizable residues D, E, K or R made at the buried positions of U1A. As expected, both deep learning (DL) and transformer (TR) models accurately predict the structure of the wild-type proteins (0 substitutions). However, as ionizable residues are progressively introduced at buried positions, up to 5-6 substitutions, both the DL- and TR-models continue to predict structures in which these residues remain buried in the core of U1A (Figure 3 **Panels A-D**), regardless of whether they are acidic (D or E) or basic (K or R). This is evident in the progressive decrease in fraction exposed as the number of substitutions increases (suggesting that they remain buried in the protein core), and in low values of structural descriptors such as RMSD and Rg (Figure 3 **Panels E-L**), indicating a high degree of similarity to the wild-type structure. Importantly, the pLDDT score remains high (above 90%), indicating strong confidence in the predicted structures (Figure 3 **Panels M-P**). As the number of substitutions increases beyond 5-6, the differences in trends between the DL and TR models become apparent. After ∼5-6 substitutions, the TR models start to recognize the incompatibility of an increased number of ionizable residues at buried positions, leading to predictions that expose the ionizable residue at the expense of generating highly open conformations with high RMSD and Rg (Figure 3 **Panels E-L**). Concomitant decrease in pLDDT score after 5-6 substitutions is also notable (Figure 3 **Panels M-P**). In contrast, DL models, such as AlphaFold2, fail to recognize that placing a large number of ionizable residues buried in the protein core violates established physico-chemical principles of protein structural stability, and continue predicting structures in which all core residues of U1A are occupied by buried ionizable residues (Figure 4). This is evidenced by a continuous decrease in the fraction exposed as a function of the increased number of substitutions (Figure 3 **Panels A-D**), and invariant RMSD and Rg for acidic residues D and E (blue or green symbols in Figure 3 **Panels E,F,I,J**), suggesting that the predicted structures are very similar to that of the wild-type protein. The pLDDT score for acidic residues (D and E) also does not change as a function of the number of substitutions (blue or green symbols in Figure 3 **Panels M-N**). In contrast, there is a small increase in RMSD (red symbols in Figure 3 **Panels G,H**) and a concomitant drop in pLDDT (from 95% to ∼80%) for structural predictions involving basic residues K and R (red symbols in Figure 3 **Panels O,P**). These changes result from differences in the sizes of the side chains upon substitution. On average, the substitutions to D lead to a decrease in the volume, substitutions to E on average lead to no volume changes, with substitutions to K and more so to R leading to an increase in sidechain volume (see **Table S2**), which, in order to be accommodated in the protein core, will require movement of the backbone. Indeed, the distance between helix α1 and sheet β2 increases by ∼6-8Å in AlphaFold2 structures with all-K or all-R substitutions (red symbols in **Figure S2**). This also allows partial exposure of some of these long and flexible side chains.

**Figure 3.**
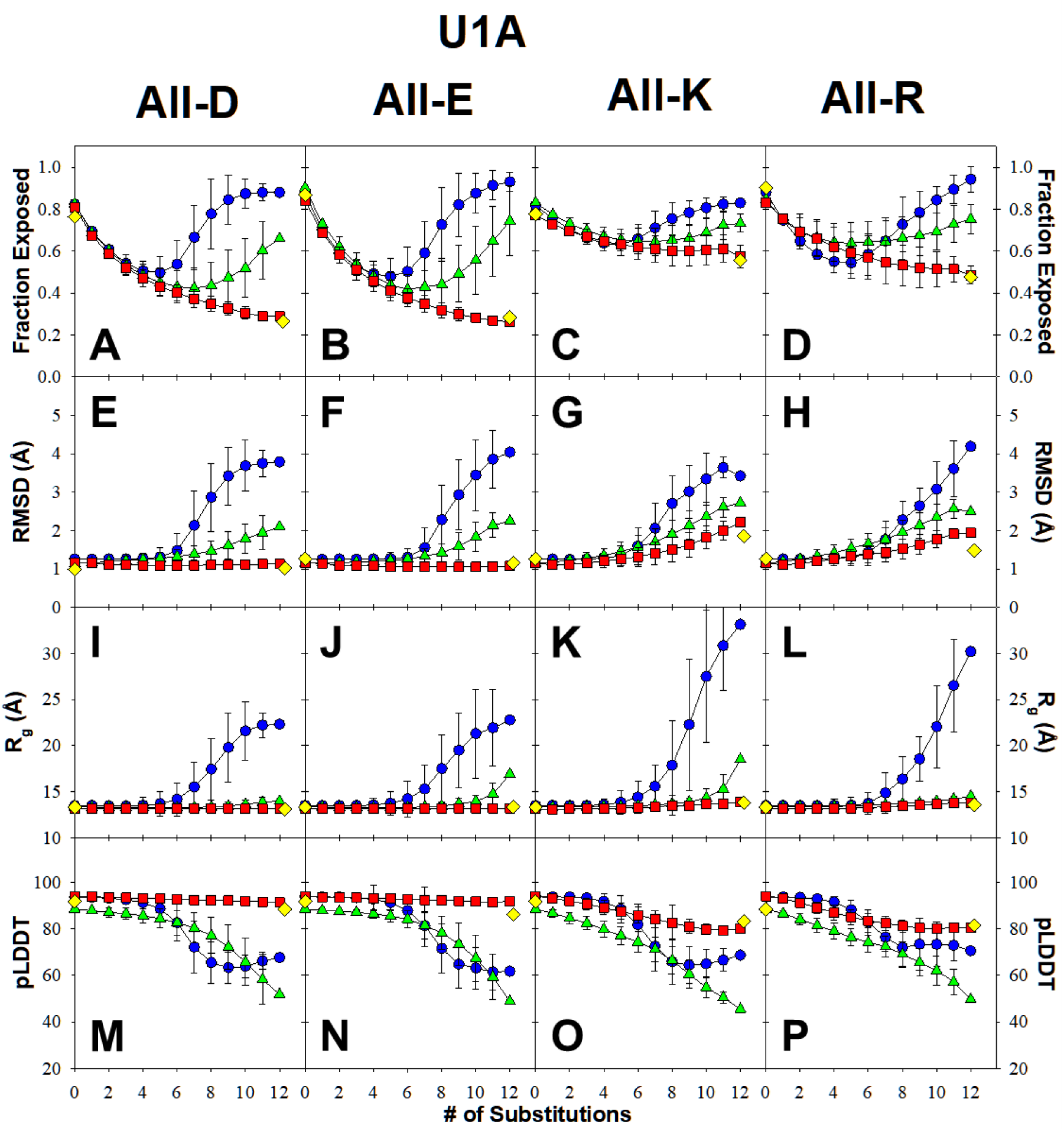
Characterization of structural models predicted by different AI models for U1A. Average fraction exposed upon substitutions (row 1, Panels A-D), average root mean square deviation from the experimental PDB structure (row 2, Panels E-H), average radii of gyration (row 3, Panels I-L), and average pLDDT score (row 4, Panels M-P) for Asp (column 1, Panels A, E, I, M), Glu (column 2, Panels B, F, J, N), Lys (column 3, Panels C, G, K, O) or Arg (column 4, Panels D, H, L, P) plotted as a function of number of substitutions. Error bars are the standard deviations of the mean calculated for all structures with the same number of substitutions. Red Squares 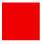 – AlphaFold2; Blue Circles 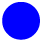– OmegaFold; Green Triangles 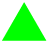 – ESMFold; Yellow Diamonds 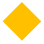 – RoseTTAFold2. See supplementary **Figure S2** for all individual data points.

**Figure 4.**
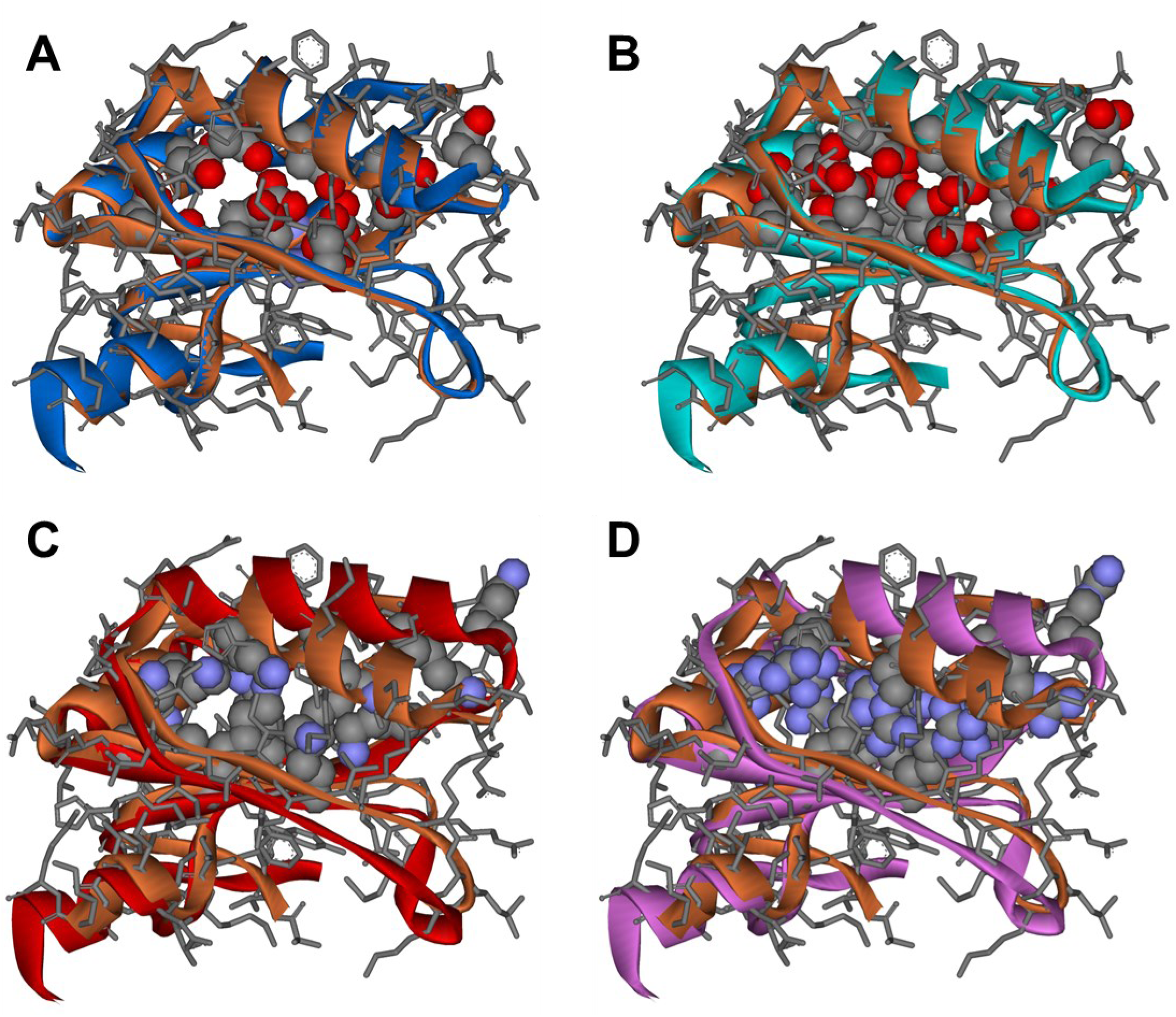
A cartoon of an overlay of the experimental structure of U1A (PDB:1URN, orange cartoon, gray sticks for bonds) with the structure of U1A variants as predicted by AlphaFold2 with all twelve core residues replaced with: Panel A - Asp (blue cartoon); Panel B - Glu (cyan cartoon); Panel C - Lys (red cartoon); Panel D - Arg (purple cartoon). The residues in the core of the variant proteins are shown in CPK representation.

Consequently, a larger fraction of exposed basic residues (red symbols in Figure 3 **Panels C-D**) than that for acidic residues (blue or green symbols in Figure 3 **Panels A-B**) is observed. This behavior seems inherent to DL models, as RoseTTAfold2 also shows similar trends (see the yellow symbols in all panels of Figure 3).

How general are the results obtained on the U1A protein? To answer this question, we applied the same type of computational analysis to two other proteins: acylphosphatase (ACP, PDB:2ACY) and TOP7 (PDB:1QYS). These proteins were selected for several reasons. First, because they are of comparable size: U1A is 101 amino acid residues long, ACP is 98 residues long, and TOP7 is 92 residues long. Second, the three proteins have a similar number of secondary structure elements: U1A has 2 helices and 4 β-strands, while ACP and TOP7 each have 2 helices and 5 β-strands, arranged into distinct topologies. Importantly, TOP7 is the first computationally designed sequence that folds into a novel topology that has not been observed in nature. Correspondingly, no MSA is available for this protein, although the experimentally determined structure is available from PDB and has been used to train DL models. The structure of ACP contains 12 positions that form the non-polar core of the protein, while the non-polar core of TOP7 contains 15 residues (see **Table S3**). The analysis of MSA for these buried positions in ACP shows that they are largely occupied by hydrophobic residues (defined are Ala, Val, Leu, Ile, Phe, Trp, Tyr) and the fraction of ionizable residues at these positions in most cases is less than 0.1%.

Figure 5 shows the results of structure predictions by TR and DL models for the ACP sequences with ionizable residues incorporated at 12 core positions (see **Table S3** for the list of positions). Again, as in the case of U1A, it appears that both deep learning (DL) and transformer (TR)- based models produce structural sets that exhibit very different trends across the four structural predictors described. However, these differences appear to be more subtle for ACP.

**Figure 5.**
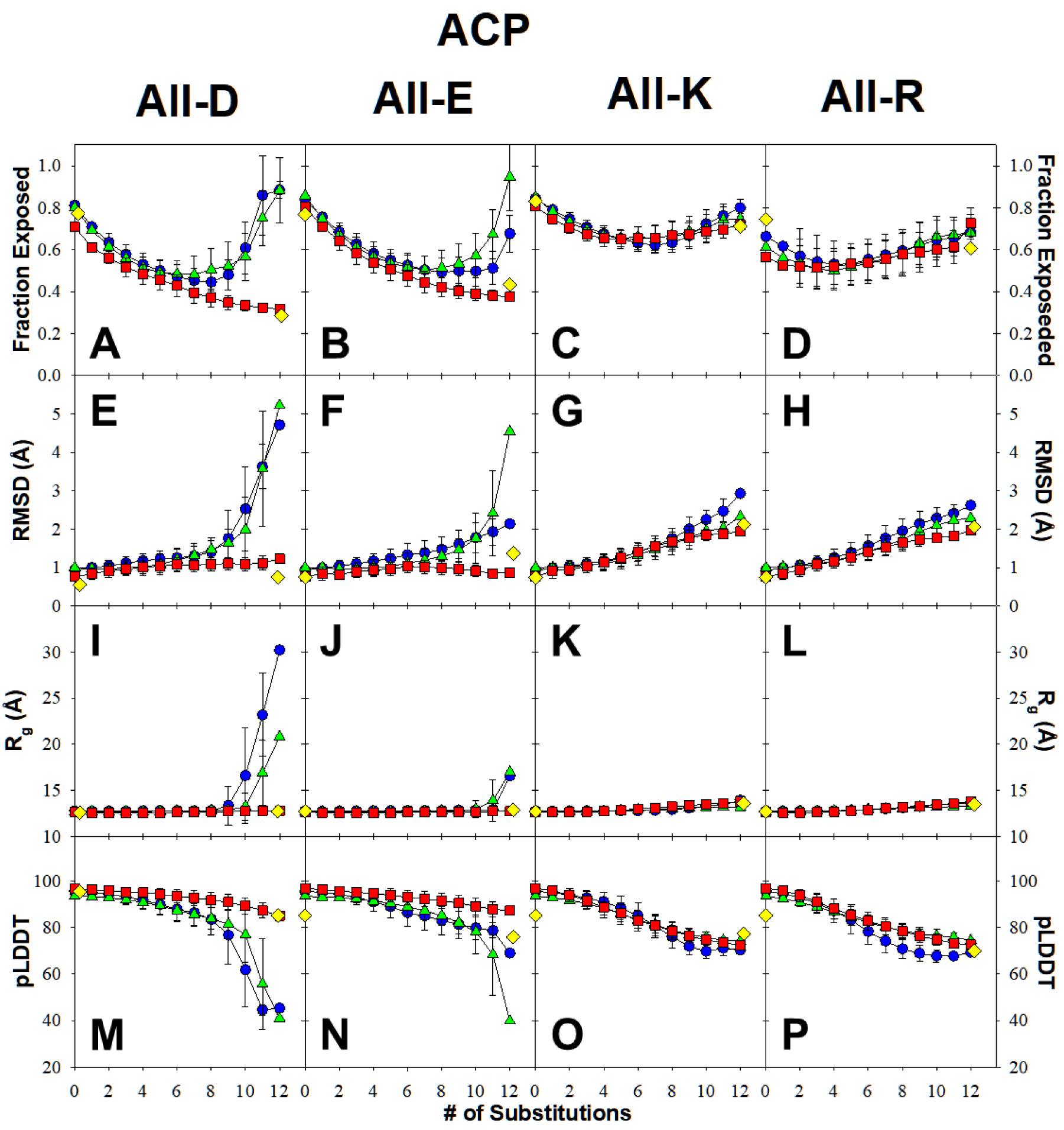
Characterization of structural models predicted by different AI models for ACP. Average fraction exposed upon substitutions (row 1, Panels A-D), average root mean square deviation from the experimental PDB structure (row 2, Panels E-H), average radii of gyration (row 3, Panels I-L), and average pLDDT score (row 4, Panels M-P) for Asp (column 1, Panels A, E, I, M), Glu (column 2, Panels B, F, J, N), Lys (column 3, Panels C, G, K, O) or Arg (column 4, Panels D, H, L, P) plotted as a function of number of substitutions. Error bars are the standard deviations of the mean calculated for all structures with the same number of substitutions. Red Squares 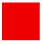 – AlphaFold2; Blue Circles 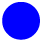 – OmegaFold; Green Triangles 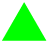 – ESMFold; Yellow Diamonds 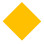 – RoseTTAFold2. See supplementary **Figure S3** for all individual data points.

TR models start to react at ∼8-10 acidic substitutions in ACP: there is a concomitant increase in the fraction exposed, RMSD, and Rg, and a decrease in the average pLDDT score (blue or green symbols in Figure 5 **Panels A-B,E-F,I-J,M-N**). For basic residues, the TR models maintain an overall topology similar to the WT and accommodate substitutions by expanding the protein core, by increasing the distance between α1 and α2. This is evident in the increase in RMSD to 3 Å, without significant changes in Rg, while the average pLDDT drops to ∼75 (blue or green symbols in Figure 5 **Panels G-H,K-L,O-P**).

For acidic residues, the DL models fail to recognize the incompatibility of the substitutions of nonpolar residues in the core of ACP with D or E, and predict a structure similar to the WT with buried ionizable residues, as evidenced by the values of RMSD, Rg, and pLDDT independent of the number of substitutions, and by low fractional exposure of the ionizable residues (red or yellow symbols in Figure 5 **Panels A-B,E-F,I-J,M-N**). For the ACP variants with the substitution to basic residues K and R, the DL models show trends similar to those of the TR models, i.e., the ionizable residues K and R have larger volumes and thus become accommodated in the core of the protein due to an increase in the distance between α1 and α2. This leads to a small increase in RMSD and unchanged Rg, because overall topology remains similar to the WT structure, while pLDDT gradually decreases to ∼75 (red or yellow symbols in Figure 5 **Panels G-H,K-L,O-P**). Overall, comparing the results for U1A and ACP, the pressure to maintain WT-like overall topology drives DL models, while TR models begin to explore conformational space that deviates from WT-like topology when faced with packing deficiencies.

The TOP7 protein has been designed de novo (47). Thus, because its sequence lacks known homologs, both DL and TR models have little sequence information, i.e., no MSA to rely on. As expected, the DL models, trained on protein structures in the PDB, including TOP7, predict compact structures with Rg values similar to those of the WT structure. The structure adjusts to accommodate substitutions in the core, and as a result, the predicted structures exhibit greater plasticity, leading to an increase in RMSD (red or yellow symbols in Figure 6 **Panels I-L**). This increase is relatively small, up to 1.7-2.0 Å for substitutions with the acidic residues D and E. However, to accommodate the larger side chains of basic amino acid residues, the predicted structure expands further than in WT, increasing the RMSD to 3-3.5 Å (red or yellow symbols in Figure 6 **Panels G-H**). The resulting fraction exposed remains relatively low (0.6), suggesting at least partial burial of the basic side chains (red or yellow symbols in Figure 6 **Panels C-D**). In all cases, the DL-predicted structures remain compact with Rg similar to that of the WT structure (red or yellow symbols in Figure 6 **Panels I-L**). Both of these might be consequences of TOP7, being a designed protein, is over-packed and strained in ways that natural proteins usually avoid (47). As a result, substitutions perturb the highly optimized packing and induce expansion of the folded structure. TR models surprisingly predict the structure of the WT sequence remarkably well, despite very few sequence fragments exhibiting homology. However, probably for the same reason, substitutions at core residues in TOP7 are predicted to diverge quickly from the WT as the number of substitutions increases. This is evidenced by a rapid increase in fraction exposed, RMSD, and Rg, with a concomitant decrease in pLDDT value to below 60 (blue or green symbols in Figure 6).

**Figure 6.**
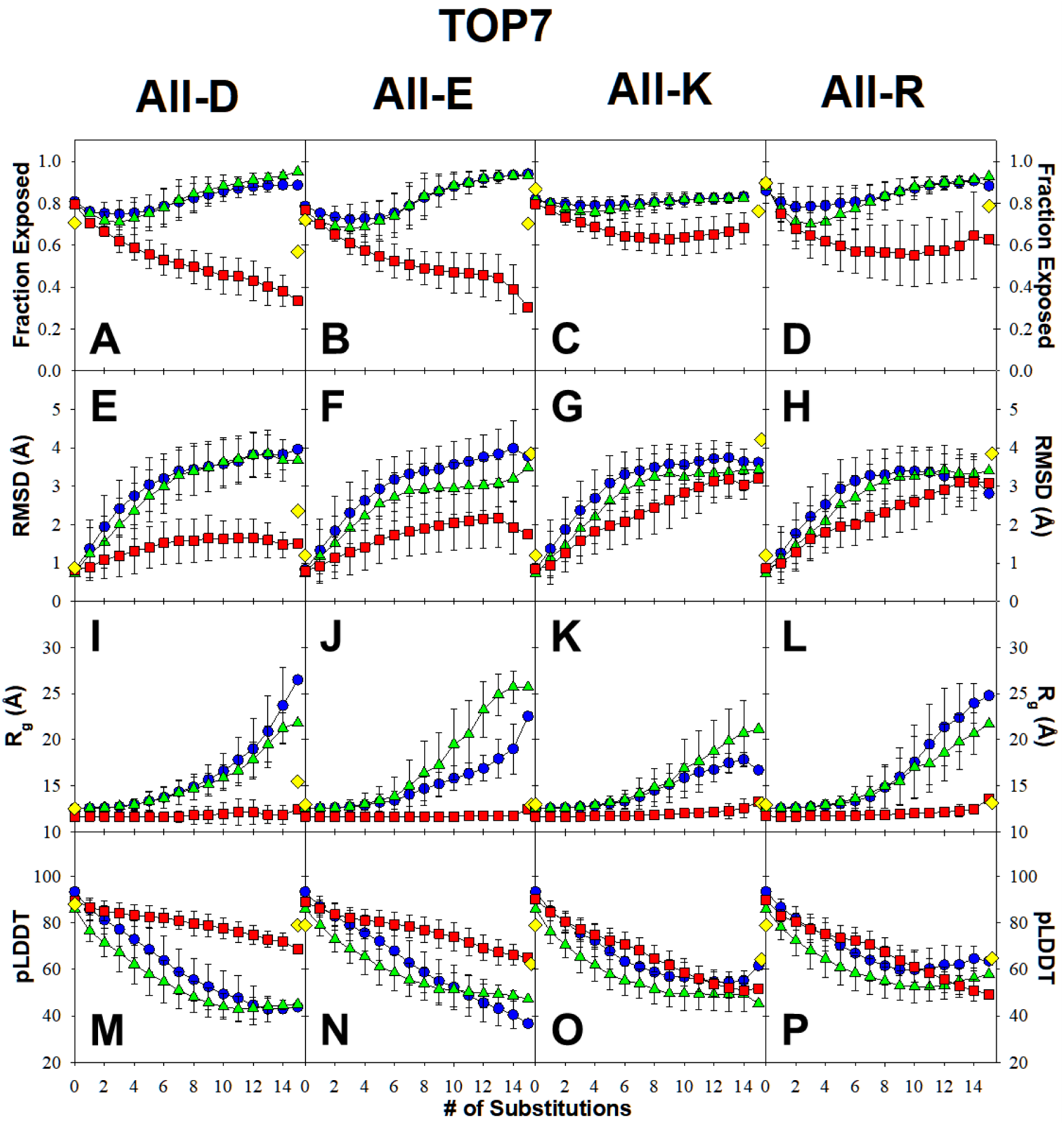
Characterization of structural models predicted by different AI models for TOP7. Average fraction exposed upon substitutions (row 1, Panels A-D), average root mean square deviation from the experimental PDB structure (row 2, Panels E-H), average radii of gyration (row 3, Panels I-L), and average pLDDT score (row 4, Panels M-P) for Asp (column 1, Panels A, E, I, M), Glu (column 2, Panels B, F, J, N), Lys (column 3, Panels C, G, K, O) or Arg (column 4, Panels D, H, L, P) plotted as a function of number of substitutions. Error bars are the standard deviations of the mean calculated for all structures with the same number of substitutions. Red Squares 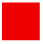 – AlphaFold2; Blue Circles 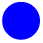 – OmegaFold; Green Triangles 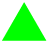 – ESMFold; Yellow Diamonds 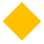 – RoseTTAFold2. See supplementary **Figure S4** for all individual data points.

Predictions by DL models, AlphaFold2 and RoseTTAFold2, are dominated by knowledge of the PDB structure for the WT sequence, with the MSA probably playing a secondary role in this context because it cannot drive exploration of alternative structures due to the absence of signals for ionizable residues at buried positions. When substitutions have larger volumes and cannot be readily accommodated in the protein core due to clashes, small structural rearrangements are introduced while preserving the overall topology. In none of the studied cases did we observe that DL-based models predicted a disordered structure, e.g. characterized by a significant increase in Rg or RMSD.

The TR models, OmegaFold and ESMFold, although not explicitly trained on structure, do have some knowledge of the known structure-sequence relationship, particularly for U1A and ACP, for which many structures from a diverse set of organisms are available in the PDB. This information is “memorized” during training and drives the prediction that sequences with a relatively small number of substitutions in the core residues of U1A and ACP have structures similar to those of corresponding wild-type proteins. Once a significant number of substitutions that disrupt the “learned” sequence patterns are introduced, TR models begin to drive structure predictions away from the WT. When sequence homology is unavailable (as in the case of TOP7), the TR model quickly recognizes interruptions in the sequence motifs and begins exploring the possible structural space based solely on the sequence it is presented with. This is one reason the OmegaFold and ESMFold models are better at predicting the structures of sequences with limited sequence homology, including singletons. Finally, the results obtained for a de novo-designed protein, TOP7, call into question the ability of DL-models to evaluate and predict structures of sequences previously unseen in nature.

### Can Physics-Based Force-Fields Detect Deficiencies in Structure Predictions?

To assess the ability of physics-based force fields, such as CHARMM or AMBER, to detect violations of physico-chemical principles in AI-based predictions, we performed molecular dynamics simulations starting from AlphaFold2-predicted structures of core variants of U1A, ACP, and TOP7 with substitutions with ionizable residues. The results were analyzed using two structural descriptors: the fraction of exposed ionizable residues and the root-mean-square deviation, RMSD, from the experimentally determined structure for the wild-type protein (Figure 7). For comparison, we also performed MD simulations starting from the wild-type protein, as reported in the experimental PDB structure (Figure 7, black symbols) as well as 3D coordinates from AlphaFold2 prediction (Figure 7, red symbols). As expected, the trajectories for the wild-type, independent of the starting structures, sample a structural ensemble with highly exposed ionizable residues and low RMSD values. In contrast, trajectories for the studied variants start with relatively buried ionizable side chains and low RMSD, but very quickly (<1 nsec) adopt conformations with highly solvent-exposed ionizable residues, enabled by large conformational changes that lead to the loss of the native-like structure, as evidenced by a significant increase in RMSD (Figure 7). Importantly, this observation is independent of the choice of force field (see Figure S3 for results from simulations using AMBER99sb-ildn).

**Figure 7.**
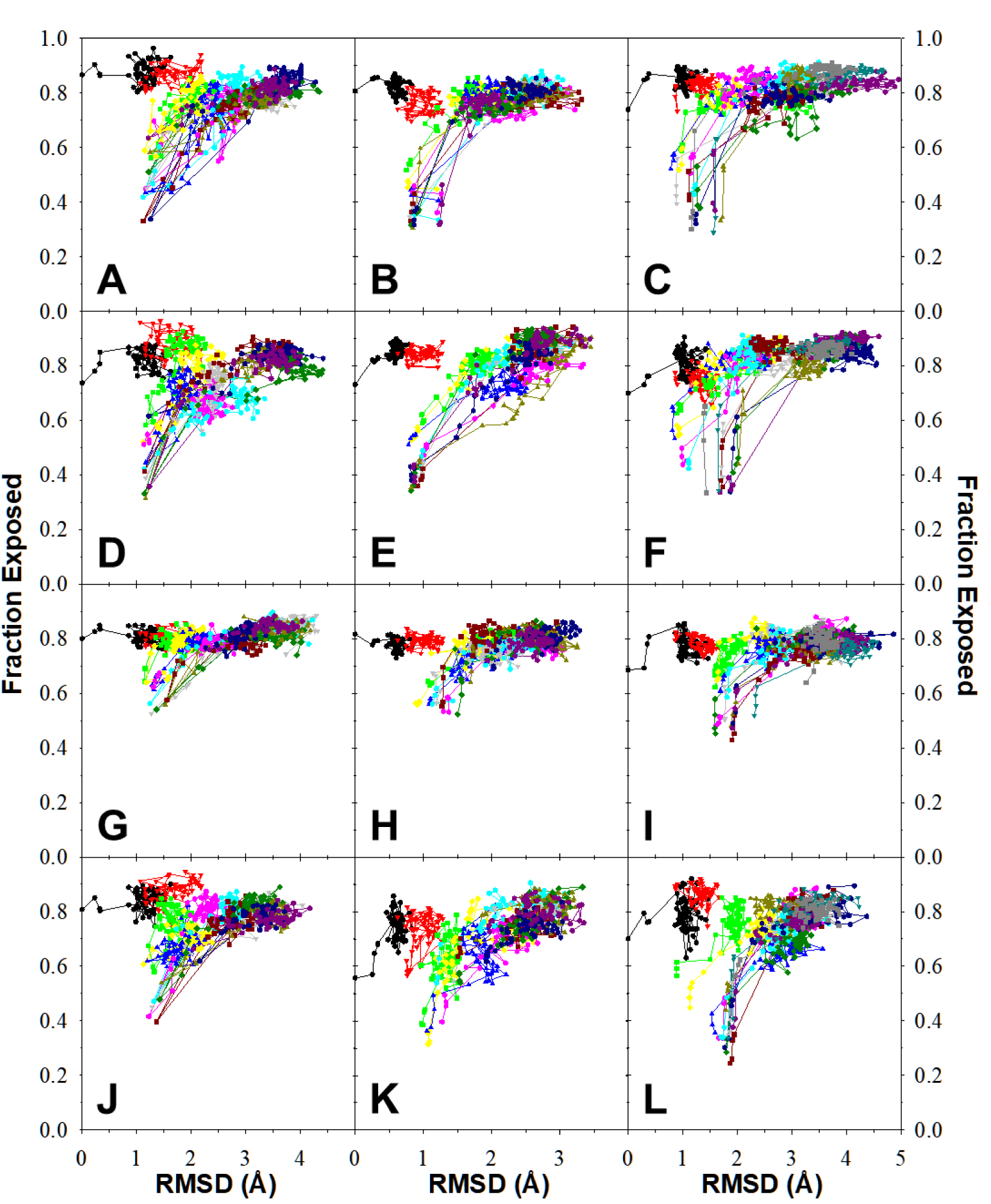
**Molecular dynamics simulations of AlphaFold2 predicted structures**. The average fraction exposed vs root mean square deviation from the experimental pdb structure during the MD equilibration (all-atom explicit solvent) in charmm36 force-field, starting from the PDB structure of the wild-type (black symbols), the AlphaFold2-predicted structure for the wild-type (red symbols), and representative sets with the increasing number of substitutions (light green 1 substitution, dark purple 12 substitutions). The first data point is the starting structure, the second is after NVT equilibration, the third is after NPT equilibration, and the remaining are successive frames collected every 50 ps, up to a total of 5 ns. Data for U1A is shown in column 1 of plots (Panels A, D, G, J), ACP column 2 of plots (Panels B, E, H, K), and TOP7 column 3 of plots (Panels C, F, I, L). See supplementary **Figure S5** for similar plots for MD equilibration (all-atom explicit solvent) in amber99sb-ildn force-field.

### Concluding Remarks

We have shown that current AI-based structure prediction methods are trained on naturally occurring protein sequences and thus cannot predict structures for sequences that introduce ionizable residues at buried positions. It appears that the “positive signal” from an observed sequence motif dominates the outcome of structure prediction. This signal does not change significantly when a rare or previously unseen amino acid substitution is introduced in this sequence fragment, and thus the assumption is made that the residue is still compatible with the “typical” structure for the parent sequence. In cases when a substitution, in addition to being rare or unseen, also introduces vastly different physico-chemical properties such as polarity, it can render the structure unstable, and thus the predicted structure will not be realized in practice. Such a sequence will either remain unstructured or, in very specific and probably rare cases, will find an alternative structure.

Overall, the TR-based models perform better than the DL models for a large number of substitutions, but they also fail to account for the unfavorable and destabilizing effects of burying ionizable residues in the hydrophobic core of the protein when presented with a few of them. It appears that one possible approach to validate AI-based predictions is to run a short MD simulation (all-atom, explicit solvent) of the predicted 3D structures for ∼50-100 ns and analyze the RMSD relative to the starting structures. If there is a drift of RMSD to higher values (>2.5Å), it will be highly probable that the predicted structures violate certain physics-based principles and should be re-evaluated.

## Materials and Methods

### Proteins: Cloning, Expression, Purification, and Characterization

The U1A expression constructs were cloned into the pGia expression vector under the control of the T7 promoter (48). Proteins were expressed and purified as described previously (33). Protein identity was validated by measuring molecular mass using MALDI-TOF. The protein concentration was measured spectrophotometrically using the extinction coefficient of U1A proteins of 5,120 a.u. for 1 M solution at 280 nm. Sequence of the wild-type protein is MAVPETRPNHTIYINNLNEKIKKDELKKSLYAIFSQFGQILDILVSRSLKMRGQAFVIFKEVSSATNALRSMQGFPFYDKPMRIQYAKTDSDIIAKMKGTF.

### Circular Dichroism (CD) Spectroscopy

All CD measurements were carried out on a JASCO J-715 spectropolarimeter (33). Far-and near-UV spectra were acquired in triplicate at 20 °C using quartz 1 and 10 mm thermostated cylindrical cuvettes, respectively. Protein concentrations were 0.2 mg/mL for far-UV spectra and 1-2 mg/mL for near-UV spectra in 5 mM sodium acetate (pH 5.5).

### Analytical Ultracentrifugation

Sedimentation equilibrium experiments were performed on a Beckman XLA analytical ultracentrifuge. Absorbance was monitored at 280 nm in long-path cells, and samples were allowed to equilibrate at three rotor speeds. Analysis of the analytical ultracentrifugation profiles obtained at speeds was done globally as previously described (49):

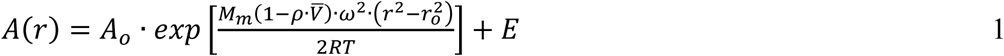

where *A(r)* is the absorbance at radial position *r*, *M_m_* is the molecular mass of the species in the cell, *V̅* is the partial specific volume of the protein, ρ is the solution density (1 g/ml), *A_o_* is the absorbance at a reference radius *r_o_*, *ω* is the rotor angular velocity, and *E* is the baseline offset term. *V̅* was calculated from the amino acid composition as described (50).

### NMR Methods

NMR spectra were acquired at 298K using a 600MHz 4 channel Bruker NEO NMR spectrometer equipped with a cryogenically cooled probe with z-axis gradients. The software package Topspin 4.4.0 was used to acquire and process data and analyze the spectra. One-dimensional ^1^H spectra were acquired using 16 scans, an acquisition time of 1.7s, and employed excitation sculpting (51) for water suppression. Two-dimensional ^1^H-^13^C one-bond correlated spectra were acquired using a SOFAST-HMQC scheme (52) with 64 scans recorded for each time increment, a recycle delay of 300 ms, and evolution times of 21 and 50 ms in the ^13^C and ^1^H dimensions, respectively.

### Analysis of the U1A Sequences

The sequence homologs of the human U1A were searched using protein Blast (https://blast.ncbi.nlm.nih.gov/Blast.cgi) with default settings, and the top 5,000 sequences (fasta aligned) were downloaded. MSA was generated using Muscle5 (5.1.linux64) using the Super5 algorithm (53) or MAFFT (v. 7.505 (54)). The frequencies of individual amino acid residues in the alignments were calculated using the consensus script (https://github.com/msternke/protein-consensus-sequence) described in the reference (55).

### Structure Prediction Methods

The structure prediction was done using AlphaFold2 (1), ESMFold (4) and OmegaFold (5). In addition, selected sequences were used to predict structures using ColabFold (6), AlphaFold3 (https://alphafoldserver.com/), and RoseTTAFold (3). The sequence of ACP used was AEGDTLISVDYEIFGKVQGVFFRKYTQAEGKKLGLVGWVQNTDQGTVQGQLQGPASKVRHMQEWLETKGSPKSHIDRASFHNEKVIVKLDYTDFQIVK. The sequence of TOP7 used was DIQVQVNIDDNGKNFDYTYTVTTESELQKVLNELMDYIKKQGAKRVRISITARTKKEAEKFAAILIKVFAELGYNDINVTFDGDTVTVEGQL.

### Calculations of Descriptors of the Predicted 3D Structures

The average exposed surface area per given type of ionizable side chain, Asp (D), Glu (E), Lys (K), or Arg (R), was calculated using the MSMS algorithm (56) and referenced to the area exposed in the Ala-X-Ala tripeptide ensemble (20 structures each), modelled using TraDes (57). The averaged pLDDT score for each structure is calculated as an average of the pLDDT of all CA atoms in each structure. The root mean square deviation (RMSD) between two structures was calculated after structural alignment using the TMalign algorithm (58).

### Molecular Dynamics Simulations

All-atom explicit solvent molecular dynamics simulations were carried out in GROMACS 2022.2 (59) using either charmm36 (60) or amber99sb-ildn (61) force-fields. The structure was solvated in a dodecahedron box, with dimensions such that all protein atoms are at least 10 Å deep in the box, and neutralized with 0.1M excess NaCl, followed by energy minimization for 1,000 steps. All simulations underwent 200 ps of constant-volume equilibration, 200 ps of constant-pressure equilibration, and up to 100 ns of production simulation at 300 K and 1 bar. We used the Parrinello-Rahman pressure control (62) with a 2 ps relaxation time, a compressibility of 4.6×10^-5^ atm^-1^, and v-rescale temperature coupling. The LINCS (63) and SETTLE (64) algorithms were used to constrain high-frequency bond vibrations, allowing the use of a 2 ps integration step. The electrostatic interactions were modeled using the smooth particle mesh Ewald method (65), with a 75×75×75 grid and fourth-order charge interpolation. Structures were extracted from the production trajectory (1 structure every 50 ps) as described (66).

### Other Methods

Van der Waals volumes of amino acid side chains were calculated using ProteinVolume (67, 68) using structural ensembles Ala-X-Ala (where X is an amino acid of interest) modelled using TraDes (57). The volumes obtained were as follows: Ala - 17 Å^3^; Val - 50 Å^3^; Leu - 67 Å^3^; Ile - 66 Å^3^; Met - 69 Å^3^; Phe - 88 Å^3^; Asp - 42 Å^3^; Glu - 58 Å^3^; Lys - 81 Å^3^; Arg - 97 Å^3^. The standard error was 1 Å3 in all cases.

## Acknowledgments

The author wishes to express gratitude to Drs. Samantha Strickler, Caitlyn Moustouka, and Scott A. McCallum for help with performing some experiments described in the paper. Computational resources were provided by the Center for Computational Innovations at RPI with software maintenance provided by Sean Collen. NMR Core at RPI is supported by NIST grant 60NANB22D167 and NIH grant 1S10OD030482.

