## Supplementary Figures S1-S5 and Tables S1-S3 for "Do AI Models for Protein Structure Prediction Get Electrostatics Right?"

***George I. Makhatadze****

*Department of Biological Sciences, Department of Chemistry and Chemical Biology, and Center for Biotechnology and Interdisciplinary Studies, Rensselaer Polytechnic Institute, 110 8$^{th}$ Street, Troy, NY, USA*

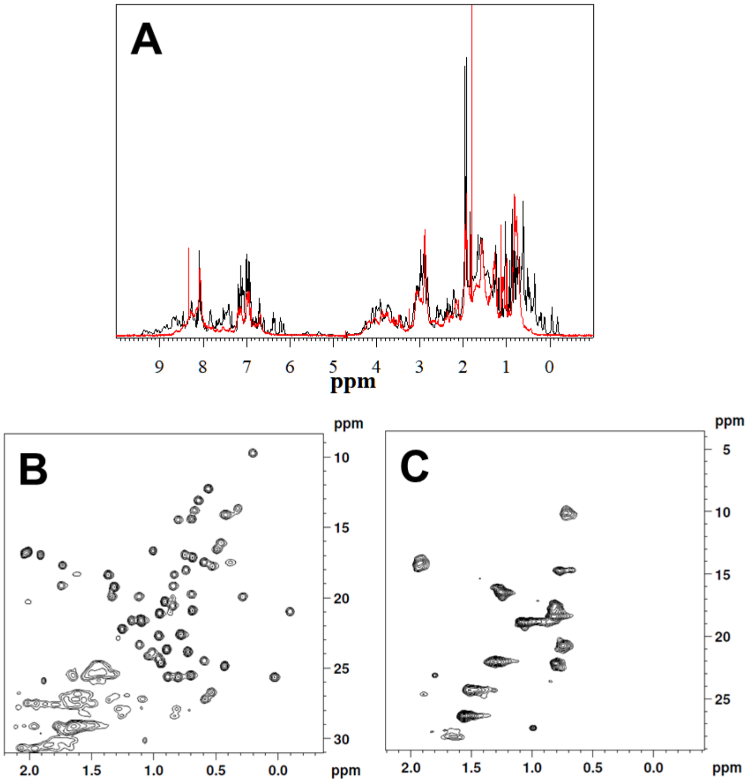


**Figure S1.** <u>**NMR-based comparison of wt-U1A and funny-U1A.**</u> **Panel A.** One-dimensional $^{1}H$ spectra comparison of wt-U1A (black) and funny-U1A (red). **Panels B & C.** Comparison of two-dimensional $^{1}H$-$^{13}C$ one-bond correlated spectra for wt-U1A (B) and funny-U1A (C). The NMR spectra of the wild-type and mutant protein constructs report a major structural rearrangement of the protein as a result of the limited modifications of the protein sequence. A high degree of chemical shift dispersion is observed in the $^{1}H$ and $^{1}H$-$^{13}C$ methyl spectra of the wt-U1A. Numerous methyl resonances are well resolved and shift substantially upfield, with several appearing below 0 ppm, and downfield amide resonances detected at approximately 9.5 ppm in proton chemical shift. These chemical shift ranges are indicative of a highly ordered structure containing a beta-sheet motif. In contrast, the spectra of the funny-U1A construct have broader resonances with few to no resolved methyl and amide peaks. These data are consistent with the CD results, in which the β-sheet content of the wild-type protein is lost, and the variant adopts largely α-helical structure (**Figure 1, Panel A**). In addition, the variant construct exhibits a longer rotational correlation time than the wild-type protein, consistent with the larger molecular mass observed in the AUC experiments (**Figure 1, Panels C-D**).

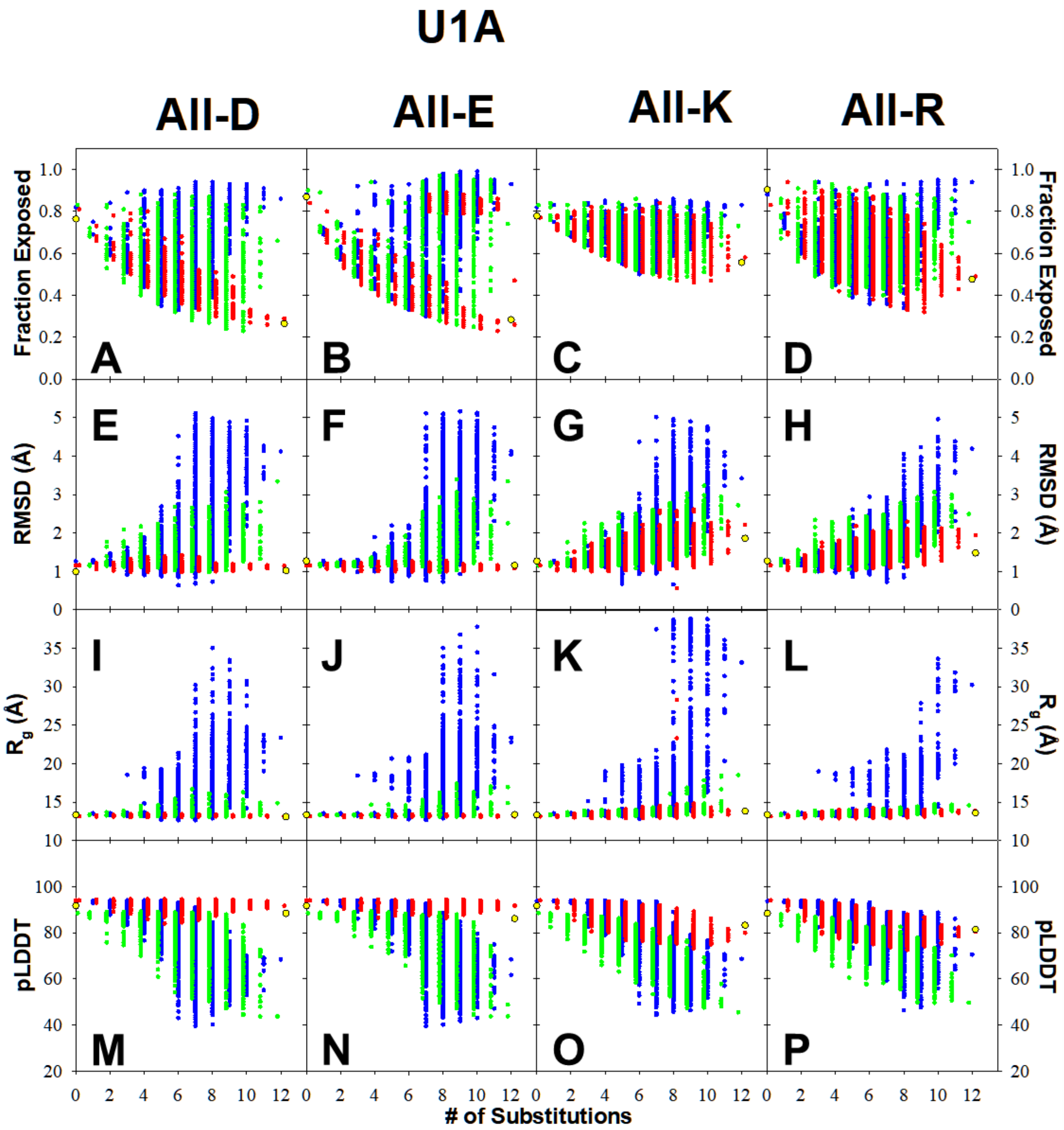


**Figure S2.** <u>**Characterization of structural models predicted by different AI models for U1A.**</u> Average fraction exposed upon substitutions (row 1, Panels A-D), average root mean square deviation from the experimental PDB structure (row 2, Panels E-H), average radii of gyration (row 3, Panels I-L), and average pLDDT score (row 4, Panels M-P) for Asp (column 1, Panels A, E, I, M), Glu (column 2, Panels B, F, J, N), Lys (column 3, Panels C, G, K, O) or Arg (column 4, Panels D, H, L, P) plotted as a function of number of substitutions. Each symbol represents an individual protein with a given set of substitutions: red – AlphaFold2; blue – OmegaFold; green – ESMFold; yellow – RoseTTAFold2. The values averaged for the given number of substitutions shown on this plot are plotted in Figure 3.

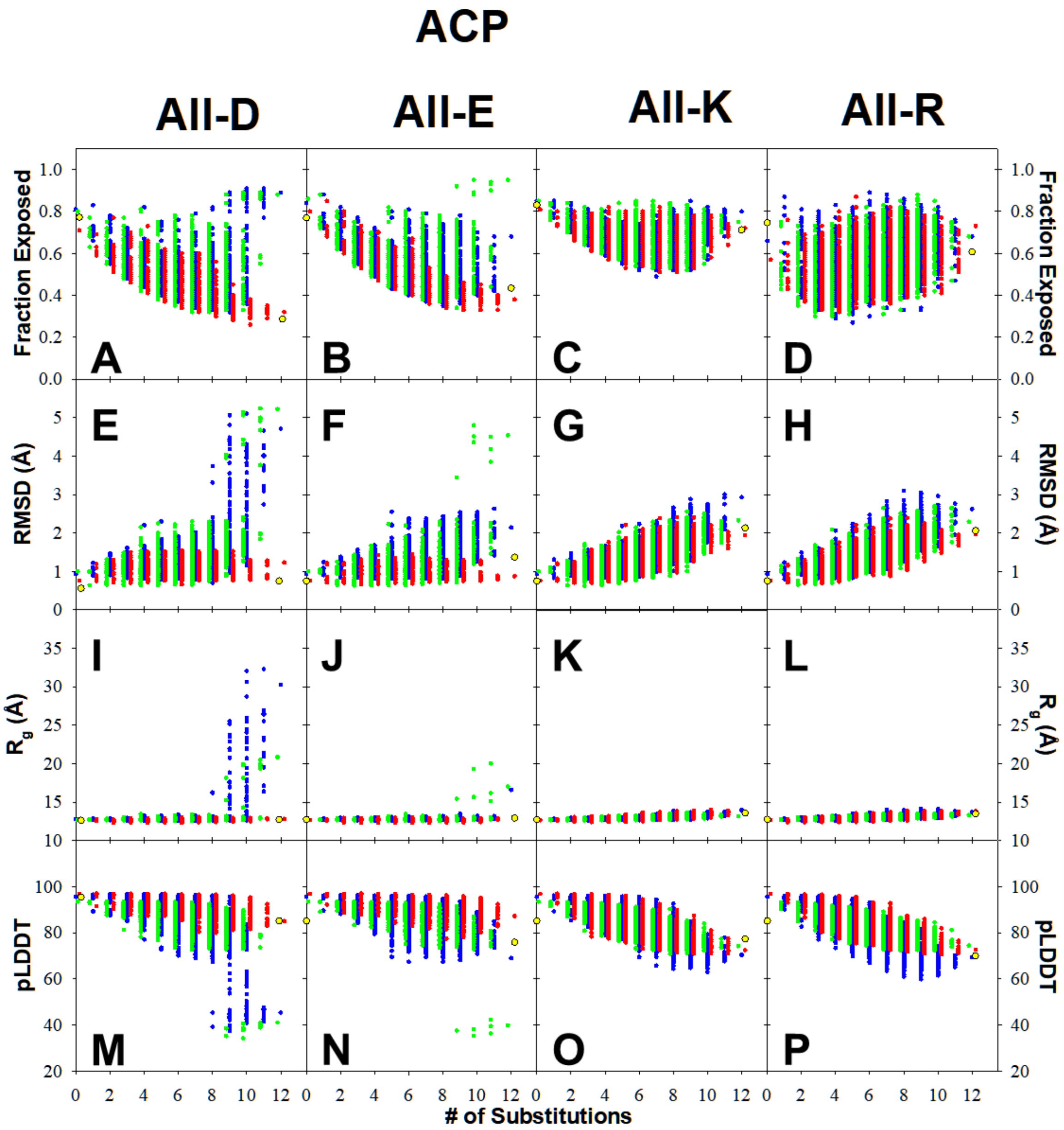


**Figure S3.** <u>**Characterization of structural models predicted by different AI models for ACP.**</u> Average fraction exposed upon substitutions (row 1, Panels A-D), average root mean square deviation from the experimental PDB structure (row 2, Panels E-H), average radii of gyration (row 3, Panels I-L), and average pLDDT score (row 4, Panels M-P) for Asp (column 1, Panels A, E, I, M), Glu (column 2, Panels B, F, J, N), Lys (column 3, Panels C, G, K, O) or Arg (column 4, Panels D, H, L, P) plotted as a function of number of substitutions. Each symbol represents an individual protein with a given set of substitutions: red – AlphaFold2; blue – OmegaFold; green – ESMFold; yellow – RoseTTAFold2. The values averaged for the given number of substitutions shown on this plot are plotted in Figure 5.

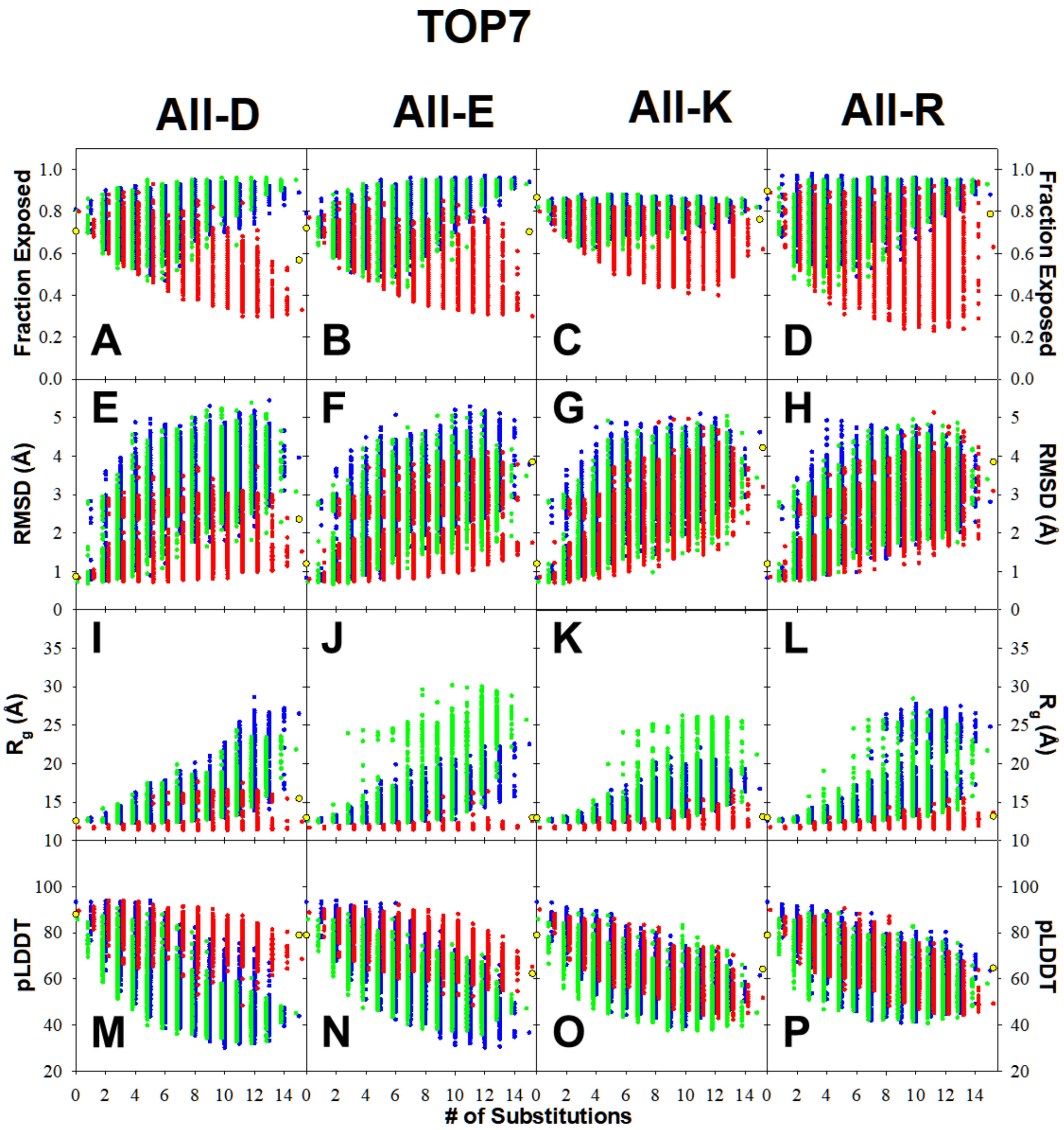


**Figure S4.** **Characterization of structural models predicted by different AI models for TOP7.** Average fraction exposed upon substitutions (row 1, Panels A-D), average root mean square deviation from the experimental PDB structure (row 2, Panels E-H), average radii of gyration (row 3, Panels I-L), and average pLDDT score (row 4, Panels M-P) for Asp (column 1, Panels A, E, I, M), Glu (column 2, Panels B, F, J, N), Lys (column 3, Panels C, G, K, O) or Arg (column 4, Panels D, H, L, P) plotted as a function of number of substitutions. Each symbol represents an individual protein with a given set of substitutions: red – AlphaFold2; blue – OmegaFold; green – ESMFold; yellow – RoseTTAFold2. The values averaged for the given number of substitutions shown on this plot are plotted in Figure 6.

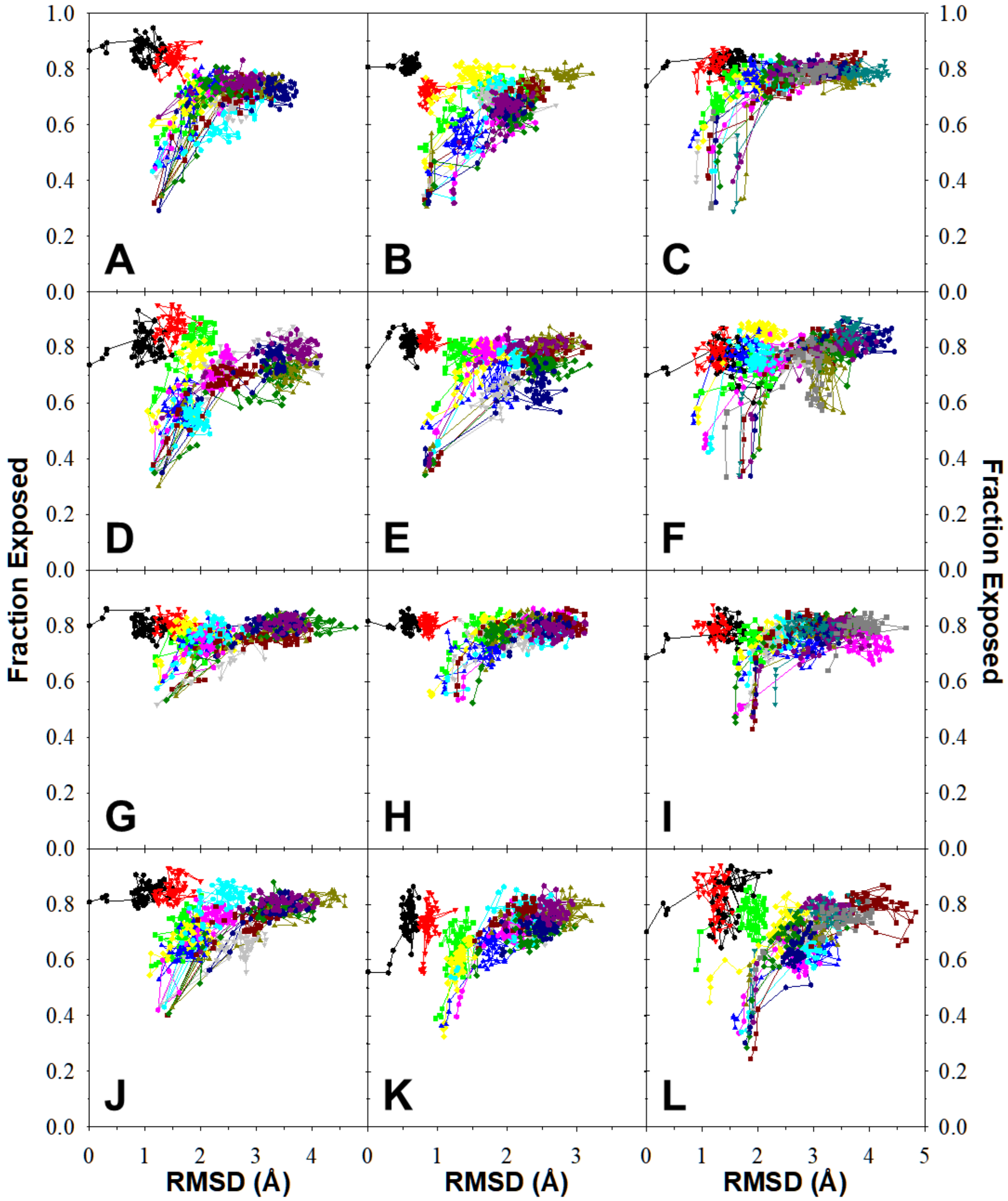


**Figure S5.** **Molecular dynamics simulations of AlphaFold2 predicted structures**. The average fraction exposed vs root mean square deviation from the experimental PDB structure during the MD equilibration (all-atom explicit solvent) using amber99sb-ildn force-field, starting from the PDB structure of the wild-type (black symbols), the AlphaFold2-predicted structure for the wild-type (red symbols), and representative sets with the increasing number of substitutions (light green 1 substitution, dark purple 12 substitutions). The first data point is the starting structure, the second is after NVT equilibration, the third is after NPT equilibration, and the remaining are successive frames collected every 50 ps, up to a total of 5 ns. Data for U1A is shown in column 1 of plots (Panels A, D, G, J), ACP column 2 of plots (Panels B, E, H, K), and TOP7 column 3 of plots (Panels C, F, I, L). Compare to **Figure 7** with similar plots for MD equilibration (all-atom explicit solvent) in the charmm36 force field.

**Table S1.** Statistical analysis of frequencies of different residues in the 5001 homologous sequences at the positions corresponding to the non-polar core of human U1A.

| Position (WT:Num) | Fraction (WT) | Fraction (Hϕ) | Fraction (ionizable) |
|---|---|---|---|
| Ile 12 | 63.40% | 99.61% | 0.04% |
| Ile 14 | 70.97% | 93.23% | 0.06% |
| Leu 17 | 83.71% | 95.62% | 0.08% |
| Ile 21 | 64.80% | 97.92% | 0.12% |
| Leu 26 | 95.79% | 97.27% | 0.04% |
| Leu 30 | 98.36% | 99.15% | 0.14% |
| Phe 34 | 96.75% | 98.06% | 0.00% |
| Ala 55 | 99.04% | 99.10% | 0.00% |
| Val 57 | 74.74% | 99.98% | 0.02% |
| Phe 59 | 95.13% | 99.88% | 0.00% |
| Ala 68 | 98.92% | 99.28% | 0.02% |
| Ile 84 | 93.75% | 99.63% | 0.08% |

For the alignment of 5,000 sequences. D, E, H, K, R are considered ionizable residues, A, V, L, I, F, P, F, Y, W are considered hydrophobic Hϕ.

**Table S2.** Volume difference introduced by incorporating ionizable residues at different positions of the non-polar core of U1A, ACP, and TOP7.

| | Volume Change (Å$^3$) Upon Substitutions With: | | | |
|---|---|---|---|---|
| **U1A** | **D** | **E** | **K** | **R** |
| **I12** | -24 | -8 | 15 | 31 |
| **I14** | -24 | -8 | 15 | 31 |
| **L17** | -25 | -9 | 15 | 31 |
| **I21** | -24 | -8 | 15 | 31 |
| **L26** | -25 | -9 | 15 | 31 |
| **L30** | -25 | -9 | 15 | 31 |
| **F34** | -46 | -30 | -7 | 9 |
| **A55** | 25 | 41 | 64 | 80 |
| **V57** | -8 | 8 | 31 | 47 |
| **F59** | -46 | -30 | -7 | 9 |
| **A68** | 25 | 41 | 64 | 80 |
| **I84** | -24 | -8 | 15 | 31 |
| **Sum of all 12:** | **-220** | **-28** | **249** | **441** |

| **ACP** | | | | |
|---|---|---|---|---|
| **I7K** | -24 | -8 | 15 | 31 |
| **V9K** | -8 | 8 | 31 | 47 |
| **I13K** | -24 | -8 | 15 | 31 |
| **V17K** | -8 | 8 | 31 | 47 |
| **F22K** | -46 | -30 | -7 | 9 |
| **L35K** | -25 | -9 | 15 | 31 |
| **V39K** | -8 | 8 | 31 | 47 |
| **V47K** | -8 | 8 | 31 | 47 |
| **L51K** | -25 | -9 | 15 | 31 |
| **V58K** | -8 | 8 | 31 | 47 |
| **M61K** | -27 | -11 | 12 | 28 |
| **L65K** | -25 | -9 | 15 | 31 |
| **Sum of all 12:** | **-235** | **-43** | **234** | **426** |

| **TOP7** | | | | |
|---|---|---|---|---|
| **I2** | -24 | -8 | 15 | 31 |
| **V4** | -8 | 8 | 31 | 47 |
| **V6** | -8 | 8 | 31 | 47 |
| **I8** | -24 | -8 | 15 | 31 |
| **L27** | -25 | -9 | 15 | 31 |
| **L34** | -25 | -9 | 15 | 31 |
| **I38** | -24 | -8 | 15 | 31 |
| **V46** | -8 | 8 | 31 | 47 |
| **I48** | -24 | -8 | 15 | 31 |

| | | | | |
|---|---|---|---|---|
| **I50** | -24 | -8 | 15 | 31 |
| **A58** | 25 | 41 | 64 | 80 |
| **L65** | -25 | -9 | 15 | 31 |
| **F69** | -46 | -30 | -7 | 9 |
| **V86** | -8 | 8 | 31 | 47 |
| **V88** | -8 | 8 | 31 | 47 |
| **Sum of all 15:** | **-255** | **-15** | **331** | **571** |

The volumes of individual side chains were obtained as described in Materials and Methods. The standard error was ~2 Å3 in all cases.

**Table S3.** Positions forming the non-polar core of ACP and TOP7

| Position ACP | Fraction (WT ACP) | Fraction (Hϕ-ACP) | Fraction (ion. ACP) | Position TOP7 |
|---|---|---|---|---|
| **I7** | 69.1% | 97.3% | 1.0% | **I2** |
| **V9** | 63.1% | 86.2% | 4.14% | **V4** |
| **I13** | 33.5% | 99.8% | 0.00% | **V6** |
| **V17** | 98.0% | 99.7% | 0.11% | **I8** |
| **F22** | 89.8% | 99.6% | 0.02% | **L27** |
| **L35** | 55.4% | 99.5% | 0.00% | **L34** |
| **V39** | 81.3% | 86.9% | 0.04% | **I38** |
| **V47** | 95.1% | 100% | 0.00% | **V46** |
| **L51** | 20.4% | 99.1% | 0.00% | **I48** |
| **V58** | 69.3% | 99.4% | 0.08% | **I50** |
| **M61** | 64.6% | 99.8% | 0.02% | **A58** |
| **L65** | 67.7% | 76.2% | 0.09% | **L65** |
| | | | | **F69** |
| | | | | **V86** |
| | | | | **V88** |

The TOP7 sequence was a de novo design, and thus, there is no MSA to analyze. D, E, H, K, R are considered ionizable residues, A, V, L, I, F, F, Y, W are considered hydrophobic Hϕ.
